# AEGIS: an annotation extraction and genomic integration resource

**DOI:** 10.64898/2025.12.04.692274

**Authors:** David Navarro-Payá, Antonio Santiago, Amandine Velt, Marco Moretto, Camille Rustenholz, José Tomás Matus

## Abstract

The GTF/GFF3 formats are the standard for storing and exchanging genome annotations. However, their flexibility often results in inconsistent and poorly formatted files across different sources, creating a major bottleneck for downstream bioinformatics analyses. Here, we present Annotation Extraction and Genomic Integration Suite (AEGIS), a comprehensive and user-friendly command-line toolkit designed to parse, validate and standardise genome annotation files. AEGIS robustly corrects common structural and formatting errors, ensuring interoperability with downstream tools. Beyond standardisation, the suite provides advanced modules for analysis, such as flexible sequence extraction (e.g. genes, CDS, proteins) with isoform handling, customisable promoter region definitions and targeted DNA motif searches. A key feature of AEGIS is its integrated workflow for comparative genomics, which combines multiple lines of evidence (i.e., sequence homology, synteny and coordinate-based lift-overs) to enable a robust gene ID correspondence and orthology assessment. We demonstrate the utility of AEGIS by comparing two major *Arabidopsis thaliana* annotations (TAIR10 vs. Araport11), successfully identifying and quantifying complex structural changes such as gene splits and fusions. AEGIS provides a unified solution for annotation quality control, feature extraction and comparative genomic analysis, simplifying complex workflows and enhancing reliability in bioinformatic research. The software is open-source, implemented in Python and is available on GitHub, PyPI, and as a Docker container to ensure accessibility and reproducibility.

## Introduction

The efficient representation and description of genomic feature annotations are fundamental requirements in modern bioinformatics. Standardised and reliable genome annotations are essential to accurately interpret gene evolution, expression, and regulation across species. While several related specifications exist, such as the more restrictive Gene Transfer Format (GTF), the General Feature Format version 3 (GFF3) has become the standard for genome annotation exchange since its release in 2004, underpinning widely used annotation pipelines, such as MAKER (Cantarel et al., 2008) and BREAKER (Gabriel et al., 2024), as well as genome browsers such as IGV (Robinson et al., 2011) and JBrowse (Buels et al., 2016). This format has been integrated into most tools requiring genome annotation data as input. It consists of nine tab-delimited columns, with the ninth column, ’attributes’, containing a semicolon-separated list of tag-value pairs. These define the relationships between features, for example, by linking exons to a parent transcript using ‘ID’ and ‘Parent’ tags in a hierarchical structure representing the complex organisation of genes and their products.

Despite the existence of a formal specification, the flexibility of the GFF3 format has led to a lack of strict adherence, which has become a significant obstacle for standardised downstream processing. Annotation files vary across organisms, frequently exhibiting inconsistencies due to the use of different source software packages or notation criteria. These discrepancies can range from simple formatting errors to more complex structural problems, hindering fundamental tasks like extracting coding sequences (CDS) for translation, calculating basic annotation statistics, or isolating promoter regions for the inspection of regulatory elements.

Common issues include missing features, incorrect or inconsistent feature coordinates, improper strand orientation and incorrectly calculated phase information (in the case of coding sequences). While certain tags like ’ID’ and ’Parent’ have predefined meanings, different annotation providers often use non-standard tags or model the same data with different notation styles. For instance, the hierarchical relationship between an exon and its parent gene can be represented differently across major databases like Ensembl, RefSeq and GENCODE, complicating the parsing process required to trace a feature’s lineage. This variability often results in files where child features reference parent IDs that do not exist, leading to parsing errors in downstream applications. Many analysis tools rely on a consistent GFF structure as input, and inconsistent or poorly formatted files can lead to a variety of errors, from outright tool failure to more subtle and potentially misleading misinterpretations of the data. Such inconsistencies force users to develop *ad hoc*, and often brittle, scripts tailored to specific file sources.

Here, we introduce AEGIS (Annotation Extraction and Genomic Integration Suite), a tool capable of parsing and standardising GFF files from various sources to ensure their interoperability and the reliability of subsequent bioinformatic analyses. AEGIS’s dual-layer design provides a powerful command-line interface (CLI) for end-users, while also allowing the use of its functionalities as a versatile Python library, making it also a foundational component for developers building custom analysis pipelines or other bioinformatics software that requires reliable handling of genomic annotations.

To showcase AEGIS’s capabilities, we have developed a range of different CLI modules for downstream analyses, including simple tasks such as calculating annotation statistics and extracting feature sequences, comparable to functionalities found in AGAT (Dainat, 2022), gffread (Pertea & Pertea, 2020), or gffutils, but also more complex comparative genomics studies. The toolkit integrates workflows for analysing gene correspondence between annotation versions in the same assembly, and for assessing orthology by combining gene model overlaps via tools like Liftoff and LiftOn with additional approaches such as reciprocal BLAST searches, OrthoFinder clustering and synteny-based analyses with MCscan, thus creating a unified solution for annotation quality control and biological inquiry.

## Implementation

The AEGIS toolkit comprises a suite of modular CLI tools designed to address distinct tasks in genome annotation processing and analysis. Its broad operating system (OS) compatibility is due to its full Python implementation. Each tool command follows the ***aegis {tool}*** format and can be called from a terminal as long as installation requirements are satisfied. All of the CLI tools make use of a set of purpose-built Python classes (Supplementary Figure S1) providing a solid framework for custom-made genome annotation functionalities as well as facilitating future AEGIS updates. To ensure the long-term reliability of these functionalities, an extensive suite of unit tests have been implemented covering core parser functions and CLI operations. This framework is integrated into a continuous integration pipeline via GitHub Actions, which validates every commit to the repository and is transparently documented in the package’s GitHub history.

While the majority of AEGIS modules such as those for file standardization (tidy, reformat), feature extraction (extract), and coordinate-based overlap analysis (overlap), are implemented as native Python functions, the orthology command requires the local installation of several external bioinformatic tools: DIAMOND for sequence homology, MCscan (from the JCVI toolkit) for synteny analysis, OrthoFinder for orthogroup inference, and Liftoff/LiftOn for annotation transfer. Due to the complex installation of these dependencies and the need for a specific fork of LiftOn to ensure full compatibility with the AEGIS workflow, we provide a pre-configured Docker image that encapsulates the entire environment, ensuring reproducibility without the need for manual dependency management. Furthermore, we have established an automated release workflow using GitHub Actions. Upon every new release, the system automatically publishes the aegis-bio package to the Python Package Index (PyPI). Simultaneously, a new Docker image is built from the repository’s Dockerfile and uploaded to both Docker Hub and the GitHub Container Registry (GHCR), ensuring that users always have access to synchronised versions across all platforms.

To systematically present the breadth of the current capabilities (Figure 1), we first describe the core utilities for foundational file manipulation, standardisation and data integration. Subsequently, we detail the modules for annotation exploration, which enable quantitative summarisation and flexible data subsetting. Finally, we present the advanced analytical modules designed for downstream biological discovery, including tools for comparative genomics and annotation comparison.

**Figure 1:**
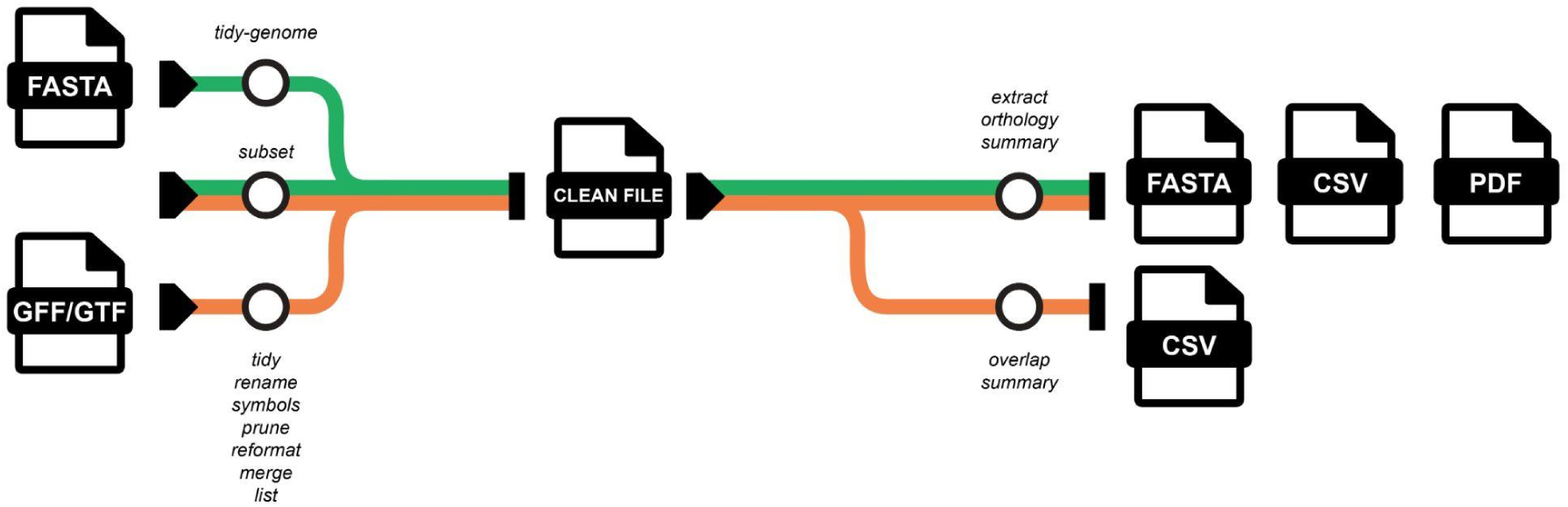
Overview of AEGIS package commands. A range of commands require annotation file(s) as input, while others simply require the assembly files. Functionalities such as sequence extraction and orthology analysis require both genome annotation and assembly file(s). The command ‘*aegis --help*’ displays a general functional description for every tool.

### Preprocessing and standardisation of genome annotation and assembly files

AEGIS has been equipped with a set of general tools to deal with common preprocessing needs and further downstream analyses. These tools provide a flexible framework for managing and standardising genome annotations for a variety of downstream applications.

The ***reformat*** tool converts annotation files between GTF and GFF formats to ensure compatibility with external software. All AEGIS tools are GTF and GFF compatible so inputs can interchangeably follow either file format (i.e. the input format is automatically determined based on file content).

The ***tidy*** command has been implemented as a standardising and error-correcting tool that can generate features if they do not exist (e.g. untranslated regions or exons). It can also polish parent-referencing, feature coordinates and phase values, while providing a set of options to customise output GFFs such as filtering for a particular biotype or whether to include untranslated regions (UTRs) in the output file. Whilst the standard exon format style for a GFF should involve the use of the ‘parent’ attributes wherever transcripts share the same exon, some GFF styles add all exon entries below each transcript regardless of repetition. AEGIS can accept both input formats and the ***tidy*** command can be used to choose which output style is chosen for exons, in order to fulfil external tool requirements. Similarly, parameters to choose the style of CDS entry ids are also provided.

The ***merge*** tool can combine any number of annotation files together whilst preventing id clashes and offering control over redundancy removal within the same loci through coordinate overlap parameters. Conversely, to reduce annotation size, AEGIS offers ***prune*** and ***subset***. ***Prune*** accepts lists of ids to remove, transcript or gene based, and solves derived parent and subfeature issues. ***Subset*** has been designed to help testing and debugging efforts by generating custom, lighter versions of annotation files with corresponding subsetting of their FASTA formatted genome, thus reducing unnecessary workloads.

The ***rename*** tool provides highly customisable modification of feature identifiers (gene, transcript, etc.). This includes removing or adding prefixes/suffixes, standardising transcript ID styles, or configuring new identifier structures for updated releases. In addition, the ***symbols*** tool allows users to append or replace gene symbols within an annotation file.

The ***tidy-genome*** tool provides similar functionality to ***rename*** and ***tidy***, but it is applied to genome assembly files. Non-chromosomal level scaffolds or organelles can be selectively removed. Moreover, it allows FASTA feature renaming and optional synchronisation of changes in a matching annotation file.

### Annotation summary report and targeted feature extraction

The **list** tool provides a convenient computational approach to generate tabular summaries of genes or transcripts from an annotation file, exporting the resulting datasets into TSV or CSV formats. Users can extensively customise the output tables by appending various metadata attributes, such as feature lengths, genomic coordinates, chromosome assignments, coding status, and gene symbols.

Additionally, the tool includes flexible filtering parameters to selectively retain or exclude specific feature classes, such as coding or non-coding elements, pseudogenes, and transposable elements.

The ***extract*** command can output multiple types of genomic features, including whole genes, CDSs, their translated protein sequences and putative promoter regions. The handling of RNA features takes a more sophisticated approach allowing precise export of all annotated transcripts or filtering for specific biotypes. For example, users can isolate only messenger RNAs (mRNAs) for expression quantification from poly(A)-selected libraries (e.g. to be used with pseudoaligners like Salmon or Kallisto), or they can extract specific classes of non-coding RNAs (ncRNAs), such as long non-coding RNAs (lncRNAs) or microRNAs (miRNAs), for studies focusing on non-coding gene function.

Beyond custom feature selection, users can control the inclusion of isoforms through distinct extraction modes (Figure 2). For instance, it can retrieve either all annotated variants for a given gene (“all” mode) or only a single, canonical representative, defined by default as the longest isoform (“main” mode). This is particularly useful for generating non-redundant datasets for comparative genomics or functional annotation. Furthermore, the tool provides specialised parameters to generate unique sets of protein or CDS sequences, either on a per-gene basis (to study protein diversity from a single locus) or across the entire genome (to create a non-redundant proteome database).

**Figure 2.**
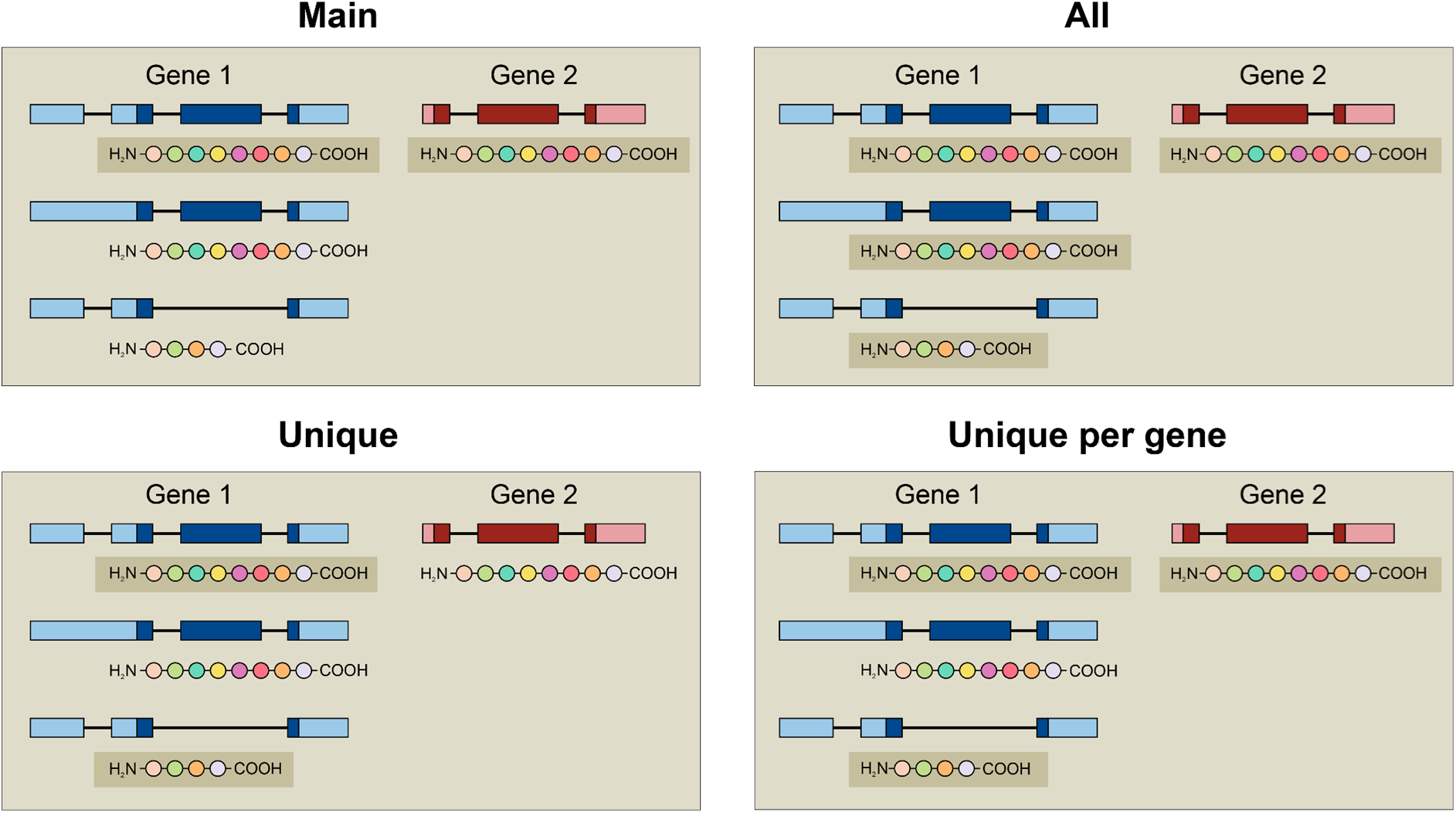
Schematic representation of the sequence extraction modes in *extract*. The figure illustrates the four modes available, via the ‘*--mode*’ argument, to control the filtering of transcript and protein sequences. The example depicts two genes: gene 1 produces two distinct protein isoforms from its three transcripts, while gene 2 produces the same protein isoform as gene 1 from one transcript. “Main” selects a primary transcript for each gene (defined by default as the one with the longest CDS); “All” extracts all annotated protein products for each gene, without any filtering; “Unique” applies a global filter, extracting only one representative unique protein sequence found across the entire dataset, effectively removing all protein redundancy within and between genes; “Unique per gene”, identifies all unique protein isoforms for each gene. Coloured boxes represent exons (CDS darker, UTRs lighter) and lines represent introns. The shaded background highlights the protein sequence output for each mode.

In the case of promoter extraction, users are not limited to a standard definition of a fixed-length region upstream of the transcription start site (TSS). The tool also allows for the extraction of regions upstream of the translation start codon (ATG), and a hybrid option which extracts the region upstream of the TSS plus the entire 5’ UTR up to the ATG (in case it is annotated). These options are particularly useful in the case of studying regulatory elements when UTRs are poorly annotated (Figure 3).

**Figure 3.**
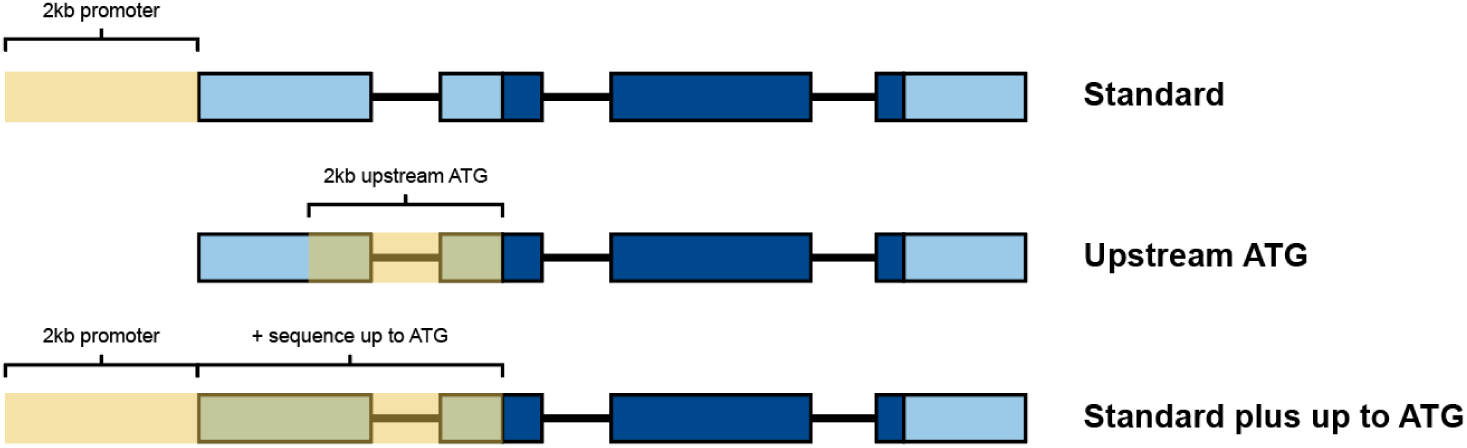
Illustration of promoter extraction strategies implemented in *extract*. Three different modes for promoter definition are supported when using the ‘*--promoter-type*’ option: (Top) “Standard”: Extracts a fixed-length region (2 kb by default) upstream of the TSS. (Middle) “Upstream ATG”: Extracts the region upstream of the ATG. (Bottom) “Standard plus up to ATG”: Combines the standard TSS-based promoter with the additional sequence between the TSS and the ATG, thereby including potential 5’ UTR elements and introns. Blue boxes represent exons (CDS darker, UTRs lighter) and lines represent introns. The highlighted yellow area depicts the extracted promoter sequence.

Finally, this tool provides extensive customisation of the output FASTA headers to facilitate data management and integration with other software. For example, users can choose to label all output sequences (e.g. transcripts, proteins) with their parent gene ID, which simplifies mapping results back to the gene level. An optional verbose output can also append genomic coordinates and other metadata directly to the sequence headers, ensuring full traceability of the extracted features.

### Comparative and functional analysis of genome annotations

#### Detecting and quantifying feature overlaps

To identify spatial overlaps between features, the ***overlap*** command includes a method where the algorithm operates in two primary modes: (1) detecting overlaps between features within a single genome annotation file (intra-annotation), and (2) comparing features between two different annotation files (inter-annotation). A fundamental prerequisite for inter-annotation comparison is that both annotation files must share the same genome assembly, ensuring a common coordinate system.

This is feasible when the annotations are different annotation versions of the same genome assembly, but for other cases, the use of external tools such as Liftoff (Shumate & Salzberg, 2021) or LiftOn (Chao et al., 2024) in advance is recommended. Alternatively, the ***orthology*** CLI described below may be more appropriate.

For each pair of potentially overlapping genes, the algorithm performs a hierarchical analysis to quantify the extent of the overlap at multiple feature levels (gene, exon and CDS). All of the calculated metrics take into account both query and target overlap percentages so that 100% is only achievable if the set of coordinates compared is exactly the same.

- Gene-level overlap: The initial overlap is calculated based on the full genomic span of the two genes (from start to end coordinates). The number of overlapping base pairs (bp) is determined and expressed as a percentage of the total length of the query gene and the target gene, respectively. The relative strand orientation (same or opposite strand) is also recorded.

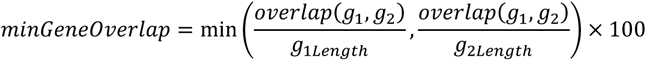

- Exon-level overlap: For a more in-depth analysis, the algorithm assesses overlap at the exon level. It compares all possible transcript pairs between the two genes. For each transcript pair, it calculates the cumulative length of all overlapping exonic regions. The maximum cumulative overlap value found among all transcript pairs is retained as the representative exon-level overlap. This value is then normalised against the total exon-spanned length of the two transcripts that produced this maximal overlap.

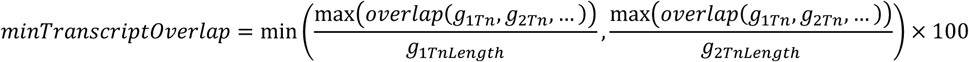

- CDS-level overlap: A similar procedure is applied to the CDSs. The algorithm identifies the transcript pair that yields the maximum cumulative overlap between their respective CDS segments. This value is subsequently expressed as a percentage of the total CDS length for each of the corresponding transcripts.

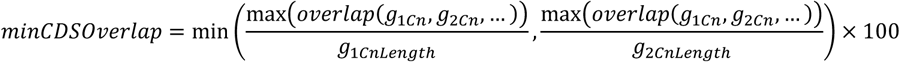

To systematically assess the structural conservation between query gene annotations and their mapped targets, we developed a custom hierarchical scoring system (Supplementary Table S1). This system quantifies the degree of overlap by prioritising the most functionally significant regions of a gene. For each query-target pair, the algorithm first attempts to calculate the minimum reciprocal overlap at CDS level. If a pair does not share overlapping CDS features (or if one of the genes is non-coding), the comparison proceeds to evaluate overlap at the exon level. If no exon overlap is found, the system finally assesses the overlap of the full gene loci (this only happens with intron-nested genes or genes with no annotated transcripts). This hierarchical approach yields a single overlap score from 0 to 11, where higher scores reflect greater structural identity (e.g. a score of 11 depicts a 100% reciprocal CDS overlap).

The algorithm also integrates pre-existing synteny information for inter-annotation comparisons, should the original annotation file, for a lifted over or transferred annotation, be provided. If a gene involved in an overlap maintains the same immediate neighbouring genes after lift-over, this status is recorded along with the overlap data. All gene overlaps can be exported in tabular format for manual inspection.

#### Orthology

Whilst ***overlap*** focuses on comparing annotation files associated with the same genome, ***orthology*** is designed to extensively compare annotations associated with different genomes. The outputs of multiple tools (internal and external) are integrated into an automated pipeline designed to generate a set of orthologues sorted into several confidence tiers.

The approach combines four complementary lines of evidence:

1. Sequence homology: the fast protein aligner DIAMOND performs reciprocal sequence searches (Buchfink et al., 2015), processing the results to identify BLAST results above a certain coverage (≥30% by default) and an e-value (≤1×10^-5^ by default) threshold. These hits are then used to determine reciprocal best hits (RBHs), reciprocal best BLAST hits (RBBHs) and the remaining results which will be only forward (fw) or reverse (rv) results depending on which annotation is considered the ‘query’. An additional identity (≥30% by default) filter for single BLAST hits (fw or rv) is also used. The threshold values are fully adjustable.
2. Synteny and collinearity: to incorporate genomic context, AEGIS uses the JCVI toolkit’s MCscan modules (Tang et al., 2024). By analysing the relative order of genes between two genomes, it identifies conserved syntenic blocks. Orthologue pairs found within these blocks receive additional support, as their relationship is preserved by both sequence similarity and genomic location.
3. Annotation lift-over and overlap analysis: AEGIS integrates annotation transfer tools like Liftoff (Shumate & Salzberg, 2021) and LiftOn (Chao et al., 2024) to project gene models from a source genome to a target genome. Uniquely, AEGIS then applies its native spatial overlap algorithm to the transferred annotations and filters overlap results using an ***overlap*** score threshold of ≥6.
4. Multi-species orthogroup inference: To incorporate phylogenetic and evolutionary information, AEGIS performs a full OrthoFinder analysis (Emms & Kelly, 2019). It extracts the primary proteome for each input species annotation, runs the OrthoFinder pipeline to cluster genes into orthogroups, and integrates the resulting pairwise orthologues from the OrthoFinder analysis into its final summary.

As output, AEGIS consolidates all sources of evidence into a single and easy-to-parse summary table with a confidence score for each putative orthologue pair. The scoring is hierarchical, where pairs classified as high confidence are primarily defined by strong evidence from lift-over based mapping, requiring a positional overlap score of ≥ 7 from either Liftoff or LiftOn (Supplementary Table S2). Pairs lacking this positional evidence are designated as medium confidence, provided they are supported by the combination of being an RBH and part of a syntenic block identified by MCscan, as well as pairs identified as RBBH alone. All other pairs identified through less stringent criteria are categorised as lower confidence, involving relationships supported by only one of the primary methods (e.g. an RBH without syntenic confirmation).

## Results

### Comparative analysis of consistency and divergence in Arabidopsis genome annotations

To showcase AEGIS’s capabilities in comparative genomics, firstly for evaluating the consistency and evolution of gene models across annotation versions within the same genome, we performed an intra-species overlap analysis between TAIR10 and Araport11 (Cheng et al., 2017; Lamesch et al., 2012). Both annotation sets are built on the latest *Arabidopsis thaliana* TAIR10 genome assembly, allowing direct comparison of feature coordinates. Using the ***overlap*** tool, we quantified the similarity between gene models in these two major *A. thaliana* annotations (Supplementary Table S3). This analysis revealed genes with conserved structural definitions across versions, as well as those showing substantial changes in their models, including structural modifications and, in some cases, changes in gene identifiers.

The initial gene-level overlap analysis revealed a high degree of concordance for the majority of Araport11 genes (31,733 with a ≥10 overlap score). This is expected for two annotations of the same genome, where genes retain their original identifier in both TAIR10 and Araport11 but where some of them may show differences in their genomic span or exonic structure. Conversely, the overlap analysis also captured cases where a single gene from the older TAIR10 annotation splits into multiple gene models in Araport11. This fragmentation often indicates a re-evaluation of gene boundaries based on improved prediction algorithms or the incorporation of additional transcriptomic evidence, where a previously annotated single locus is now recognised as two or more distinct genes. For instance, the gene ID *AT2G52240* in TAIR10 was found to have a partial overlap with *AT2G52240* and *AT2G52245* in Araport11, with the latter being a small transcript variant coming from the previous model (Figure 4A). The opposite scenario (i.e., gene merge cases), was also observed, where two or more distinct TAIR10 genes were reported to overlap significantly with a single, larger Araport11 gene (e.g. *AT1G27595*) (Figure 4B). This analysis allows users to track structural gene model changes over annotation updates, which are not always provided in such a level of detail.

**Figure 4.**
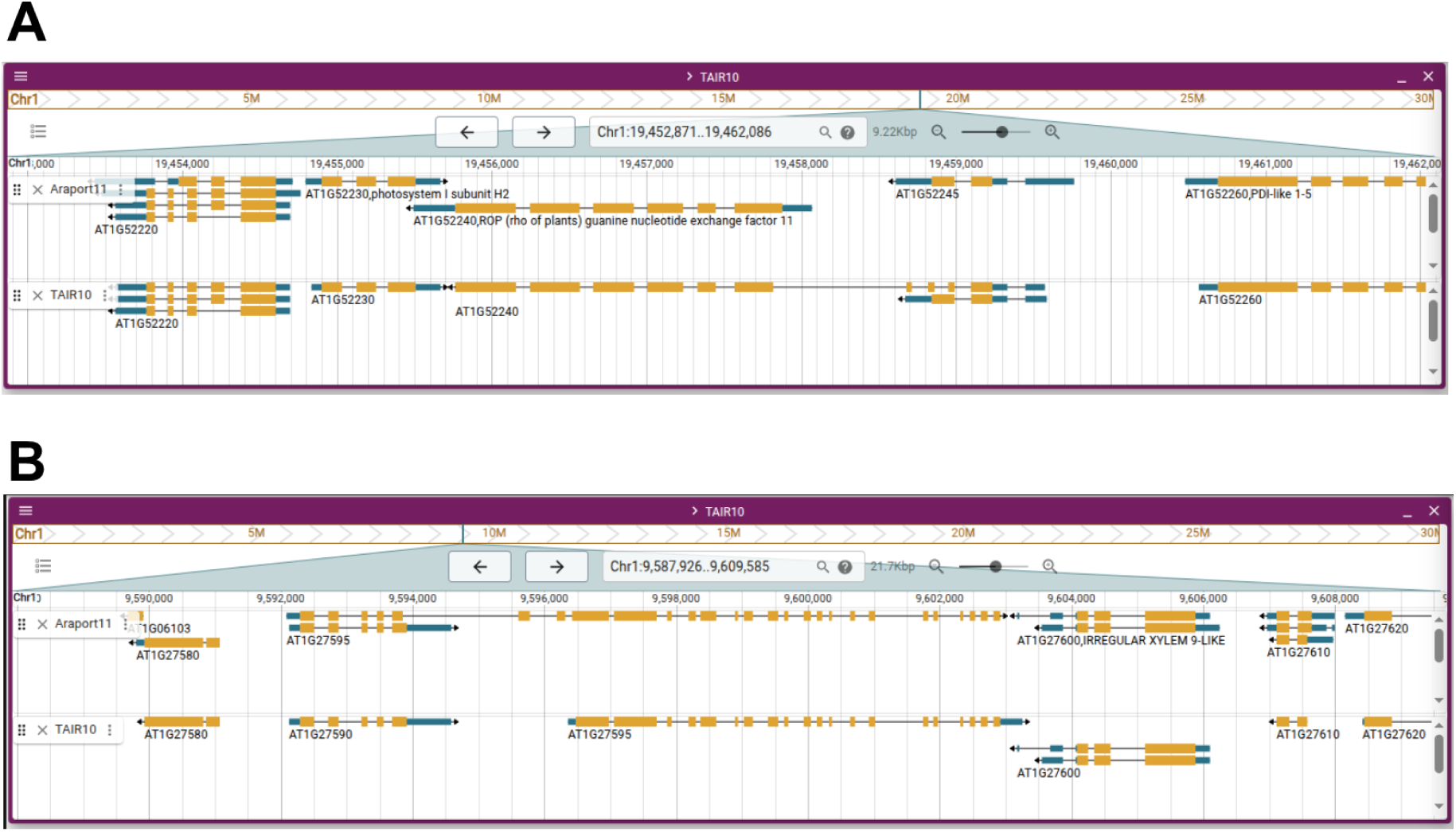
Annotation version differences as detected by *overlap*. Views of different regions in chromosome 1 in Jbrowse2, with examples of gene model changes between Araport11 (top track in each panel) and TAIR10 (bottom track) structural annotations (Arabidopsis thaliana TAIR10 genome assembly). **(A)** Example of a gene model split. The TAIR10 annotation identified a single long gene, *AT1G52240*, whereas Araport11 resolved this as two distinct genes: a redefined and shorter *AT1G52240*, and a new gene model; *AT1G52245*. **(B)** Example of a gene model merge. In the TAIR10 annotation, this locus contained two separate genes, *AT1G27590* and *AT1G27595*. The updated Araport11 annotation merged these into a single, longer gene model, *AT1G27595*. In both panels, boxes represent exons (CDSs in yellow and UTRs in blue) and black horizontal lines represent introns.

### Orthology between grapevine, tomato and Arabidopsis genomes

Furthermore, to demonstrate the utility of AEGIS in an inter-species context, we compared genome annotations from *Arabidopsis thaliana*, tomato (*Solanum lycopersicum*) and grapevine (*Vitis vinifera*). This cross-species comparison enabled the identification of conserved orthologues and highlighted structural differences in their gene models.

The annotations and assemblies of grapevine (annotation v5.1, assembly T2T.v5; Shi et al., 2023), tomato (annotation ITAG4.0, assembly ITAG4; Hosmani et al., 2019) and Arabidopsis (annotation Araport11, assembly TAIR10; Lamesch et al., 2012) were compared with ***orthology*** resulting in a list of associated gene ID pairs between annotations, supporting evidence details and a qualitative summary score. This process resulted in a comprehensive list of orthologous gene pairs, each with all of the supporting evidence details and a qualitative summary score (Figure 5A).

**Figure 5.**
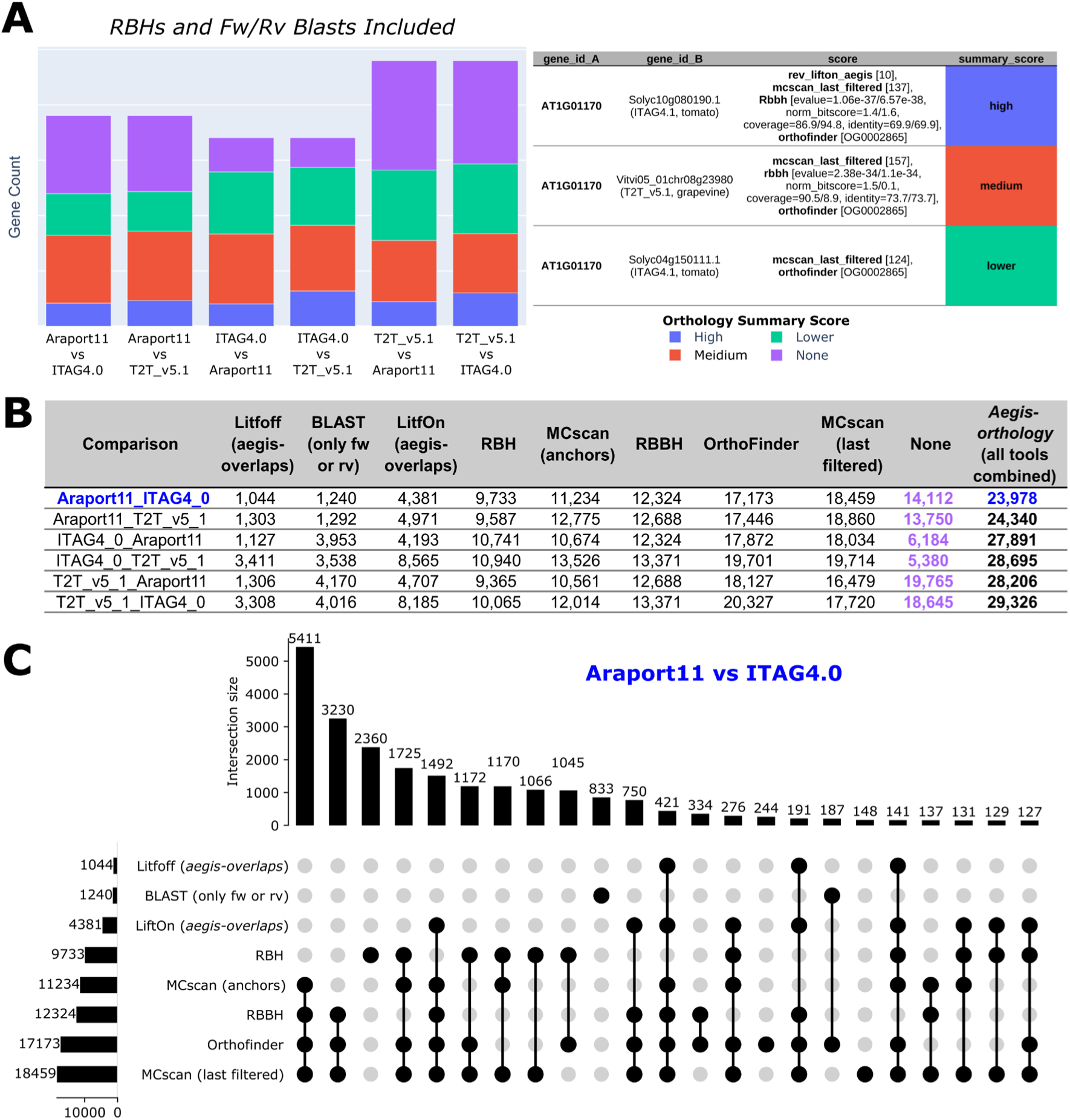
Overview and performance of the orthology tool. Used command: “*aegis orthology Araport11_GFF3_genes_transposons.20250813.gff ITAG4.0.gff3 5.1_on_T2T_ref.gff3 –annotation-names Araport11,ITAG4_0,T2T_v5_1 --group-names Arabidopsis,Tomato,Grapevine –genome-files TAIR10_genome.fasta,S_lycopersicum_chromosomes.4.00.fa,T2T_ref.fasta*”. The reported counts represent the number of genes in a query annotation that have at least one orthologue identified in the target annotation. **(A)** Summary of identified orthologues for Arabidopsis, tomato, and grapevine genes. The stacked bar chart (left) shows, for each annotation, how its genes are distributed according to whether an orthologue is present in the target species and, if so, the confidence of the best-supported orthologue. These qualitative levels are based on the combined results of individual tools, as shown for gene *AT1G01170* on the right. **(B)** Orthologue counts per tool result for each query annotation when compared to different target species. The combined results represent an improvement over individual tools in the number of genes with at least one orthologue. **(C)** UpSet plot illustrates the intersections of orthologue predictions among different methods for the Arabidopsis (Araport11) versus tomato (ITAG4.0) comparison. The vertical bars represent the number of genes with detected orthologues in each intersection (indicated by connected dots), while the horizontal bars on the left show the total number of genes with at least one orthologue identified by each method.

More than half of the genes of each genome, between the three species compared, had at least one orthologue in each pairwise comparison. For instance, 23,978 Arabidopsis genes had at least one orthologue in tomato when looking at the combined results from all tools. This combined value is higher than any of the individual metrics, supporting the benefit of the approach (Figure 5B). The Arabidopsis (Araport11) and tomato (ITAG4.0) comparison shows a degree of overlap amongst individual ***orthology*** tool results (Figure 5C). The greatest set (5,411 genes) corresponds to the shared findings of MCscan (‘anchors’ and ‘last_filtered’), RBBH and OrthoFinder, suggesting that the output of these tools is highly concordant. Nevertheless, individual tool results also manage to uniquely detect orthologues for a considerable number of genes (e.g. 2,360 genes are singularly detected by RBHs), demonstrating that each tool can contribute to the final combined result.

The output table with the default ***orthology*** command shown in Figure 5 results in over 900,000 orthologue gene-pairs. The same tool can be used with a different parameter to produce a lighter output (Supplementary Table S4). In this case, ∼125,000 orthologue gene-pairs were identified, which still manages to find orthologue matches for a sizeable portion of annotated genes (e.g. more than 50% for both Arabidopsis and tomato). This drop mainly affects the lower-confidence orthologues (Supplementary Figure S2A) and there is still a benefit in combining the different results of each tool (Supplementary Figure S2B).

### Computational performance and benchmarking

To evaluate the computational efficiency of AEGIS, we benchmarked its core utilities against comparable functionalities in an existing state-of-the-art toolkit, specifically AGAT. Benchmarks were executed on a workstation equipped with an Intel Core i9-10900 and 32 GB, recording both total execution time and memory usage (RAM).

To ensure reproducibility and consistent execution environments, all evaluated tools were containerised and executed using Docker. Upon the initialization of a tool’s container, the monitoring script continuously polled the Docker Engine API via the native docker stats command to capture real-time memory usage. Data points were recorded at a sampling interval of 0.1 seconds throughout the entire lifecycle of the containerised process. The resulting telemetry data was logged as a time-series dataset.

The comparative analysis was performed using small (*Chlamydomonas reinhardtii*), medium (*Arabidopsis thaliana*), and large genomes (*Homo sapiens*). We evaluated performance across three standard bioinformatic tasks: annotation parsing and standardisation, conversion to GTF, and protein extraction.

Across all tested benchmarks and genome sizes, AEGIS demonstrated a significantly higher computational efficiency compared to AGAT (Supplementary Figure S3). In terms of execution time, AEGIS completed the tasks of GFF tidying, GTF conversion, and protein extraction around 3-6 times faster than the corresponding AGAT scripts. This performance gap was particularly evident in the *Homo sapiens* dataset, where for instance, in GTF conversion, AEGIS finished in 194 seconds compared to AGAT’s 1,101 seconds.

Regarding memory management, although AEGIS reaches a peak memory usage more rapidly than AGAT, it maintains this footprint for a much shorter duration, drastically reducing the total computational cost. While AGAT showed a steady and prolonged memory plateau, AEGIS optimised resource release immediately upon task completion. These results indicate that AEGIS is highly scalable and particularly well-suited for large genomic annotations.

## Discussion

Implemented in Python, AEGIS provides both a powerful command-line interface for pipeline integration and an object-oriented library for custom script development, offering a flexibility absent in tools designed for one or a few tasks. It incorporates functionalities found across several tools, including robust file parsing, format conversion, filtering, and statistical reporting, matching the feature sets of mature tools like AGAT. The analysis of genome annotation files is supported by a diverse software range, which can be broadly categorised into comprehensive command-line toolkits (e.g. AGAT, GenomeTools), specialised utilities (e.g. GffCompare, TransDecoder), and programmatic libraries (e.g. gffutils). While established toolkits provide several functionalities for genome annotation handling, this new package offers an advantage in terms of parsing capabilities, being able to process complex annotation files that cause other tools to fail. For instance, AGAT successfully resolves overlapping exons within individual transcripts, but it fails to consolidate identical exons shared across multiple transcripts. Instead, AGAT redundantly duplicates these shared features, assigning each iteration a unique ID and restricting them to a single parent. In contrast, AEGIS can optionally produce a much more compact GFF3 format by making use of the multi-parent feature of the official GFF3 specification, thereby reducing unnecessary data redundancy. Moreover, some known errors in AGAT, such as the “Bio::Root::Exception” caused by long lines in FASTA files when extracting feature sequences, have been avoided when designing AEGIS. AEGIS is specifically engineered to handle such edge cases, with a codebase designed for robustness and modular reuse. Python was selected as the implementation language to maximise these attributes, favouring maintainability and flexible data handling over the performance optimisations typical of lower-level languages.

In terms of performance limitations, the AEGIS package has been benchmarked favourably against AGAT for the functionalities that were compared. Nevertheless, the performance speed of AEGIS or AGAT compared to other C/C^++^ tools such as GFF Utilities and GenomeTools is still limited based on the chosen languages themselves, Python and Perl, respectively. For basic standard operations GFF Utilities and GenomeTools may still be the option of choice, however, these offer less flexibility and robustness with respect to edge cases. Moreover, whilst AEGIS is able to deliver GFF3 recommendations more effectively in the outputs, AGAT offers greater flexibility when dealing with particular GTF format flavours.

AEGIS includes several high-level analytical capabilities not found in other examined software (Supplementary Table S5). It provides modules for identifying a gene’s main isoform, extracting a range of sequences, manipulating FASTA files together with their corresponding annotation files, detecting overlaps between features, and comprehensive orthology analysis. We showed that we were able to quantify and categorise intra-species changes between annotation versions using the ***overlap*** command. It generated a detailed and up-to-date table of gene model correspondences, extending beyond a static list of obsolete identifiers (such as the ones provided in Araport11; (Cheng et al., 2017), where users can inspect and trace gene model differences between annotation releases.

Moreover, AEGIS can be used to compare annotation files from different genome assemblies, addressing a complex comparative orthology analysis that lies beyond the scope of existing tools like GffCompare (Pertea & Pertea, 2020). While GffCompare focuses on RNA processing, comparing coordinates primarily at the intron and exon levels to match intron chains, AEGIS ***overlap*** is more focused on whether an overlapping pair of genes produce a similar functional product. It calculates mutual overlap percentages at the gene, exon, and CDS levels, specifically validating that CDS segments maintain the same reading frame and retrieving exact metrics for each pairwise correspondence. The***orthology*** tool integrates the overlaps and the output of several other tools and can be used to study genome evolution and even assist in the identification of core and dispensable genes when dealing with pangenomes. We found grapevine orthologues for 24,350 tomato genes and tomato orthologues for 24,099 grapevine genes, a substantial increase over previous studies, which reported 16,454 and 15,631 orthology relationships in tomato and grapevine, respectively (Ambrosino et al., 2018). By also providing medium and lower confidence orthologues, we highlight the ability to extract more comprehensive evolutionary signals from genome annotations, providing a stronger basis for comparative genomics and functional inference.

While the ***orthology*** example use case focuses on plant genomes, we expect that this tool and the rest of the AEGIS suite will be broadly applicable to the analysis of animal and other eukaryotic genomes as well. We anticipate that AEGIS will be a valuable framework as it unifies essential manipulation tasks in genome annotation around a flexible object-oriented core from which further functionalities can be reliably developed.

## Availability and requirements

AEGIS is available under the GPL-3 Open Source license. The version of the package associated with this work is distributed as a Python package in the Python Package Index as aegis-bio (https://pypi.org/project/aegis-bio/). The source code, along with installation and implementation instructions, is also available at GitHub (https://github.com/Tomsbiolab/aegis). A readme file is included for a comprehensive description of its features and their execution. AEGIS is also available as a containerised CLI along with other tools required for some functionalities, available as a Docker Image at Docker Hub (https://hub.docker.com/r/tomsbiolab/aegis). The package is OS independent.

## Acknowledgements

This work was supported by the grants Valinet (PID2021-128865NB-I00) and Metacell (PID2024-163039NB-I00) awarded to J.T.M. from the Ministerio de Ciencia, Innovación y Universidades (MCIU, Spain), Agencia Estatal de Investigación (AEI, Spain /10.13039/501100011033), and Fondo Europeo de Desarrollo Regional (FEDER, European Union), and to the PROMETEO grant (PROMETEO/2021/056) awarded to J.T.M. and its associated doctoral grant (PROMETEO/2021/056-01) granted to A.S., by the Generalitat Valenciana (GVA). Further support was provided by the Grapedia COST innovators grant (IG17111) awarded to J.T.M. and C.R. This research also received a grant (IEC-BG1-2023-4), awarded to J.T.M., from the Institute for Catalan Studies (IEC) within the framework of the first call for biogenome (genomics of biodiversity) research grants. Testing and debugging was carried out on the HPC cluster Garnatxa at the Institute for Integrative Systems Biology (I^2^SysBio).

## Supplementary Information

**Supplementary Figure S1:**
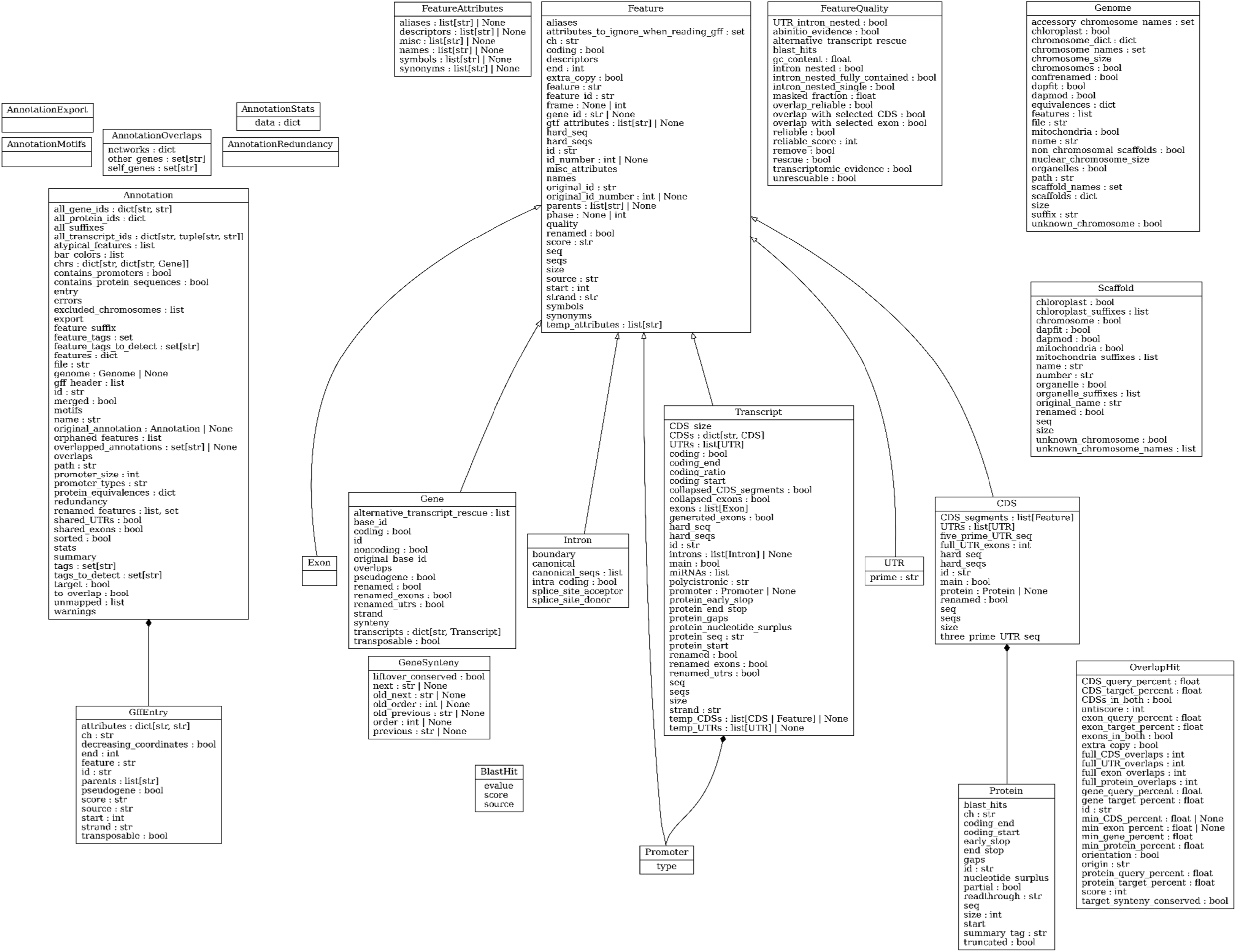
UML diagram of AEGIS Python classes and their attributes. Custom classes have been made for each annotation feature and wherever useful inheritance has been used to ensure code efficiency and reliability. All of the classes have been exposed at the top level of the package so they can be directly imported in Python, i.e. ‘*from aegis import Annotation*’ without having to know the specific module where the class is located. A detailed UML diagram including class methods can be found in the package’s main page (https://github.com/Tomsbiolab/aegis)

**Supplementary Figure S2.**
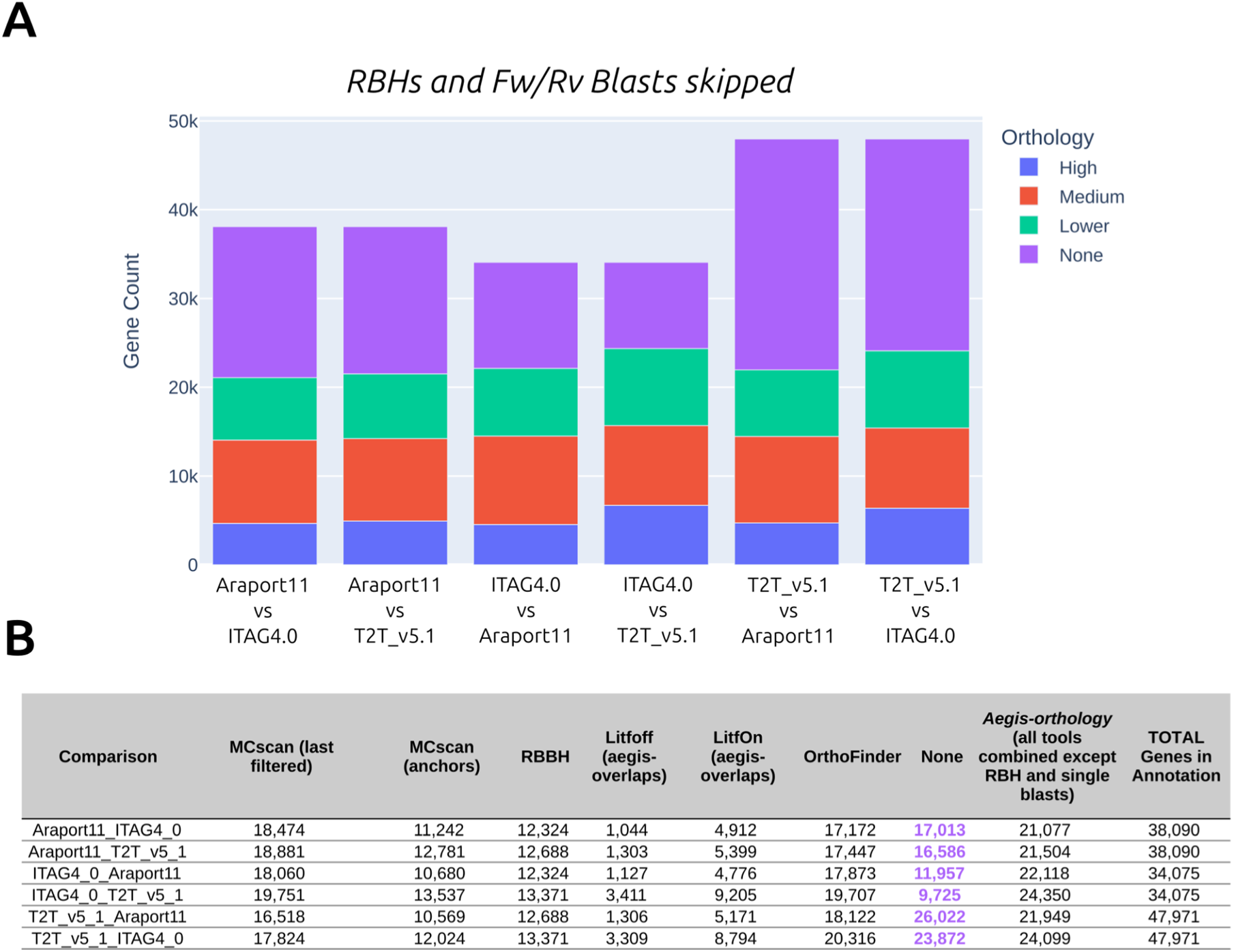
Results of *orthology* tool skipping secondary reciprocal and single forward or reverse BLAST hits. The ***orthology*** tool can be run with the ‘*--skip_rbhs*’ option to obtain a smaller output table that nevertheless still manages to detect orthologues for many genes. The exact output of this ***orthology*** is included in Supplementary Table S4. **(A)** The number of genes with no detected orthologues is greater in this case than with the default options (Figure 5A). **(B)** The combined results of ***orthology*** report more genes with at least one orthologue when compared to individual tool results.

**Supplementary Figure S3.**
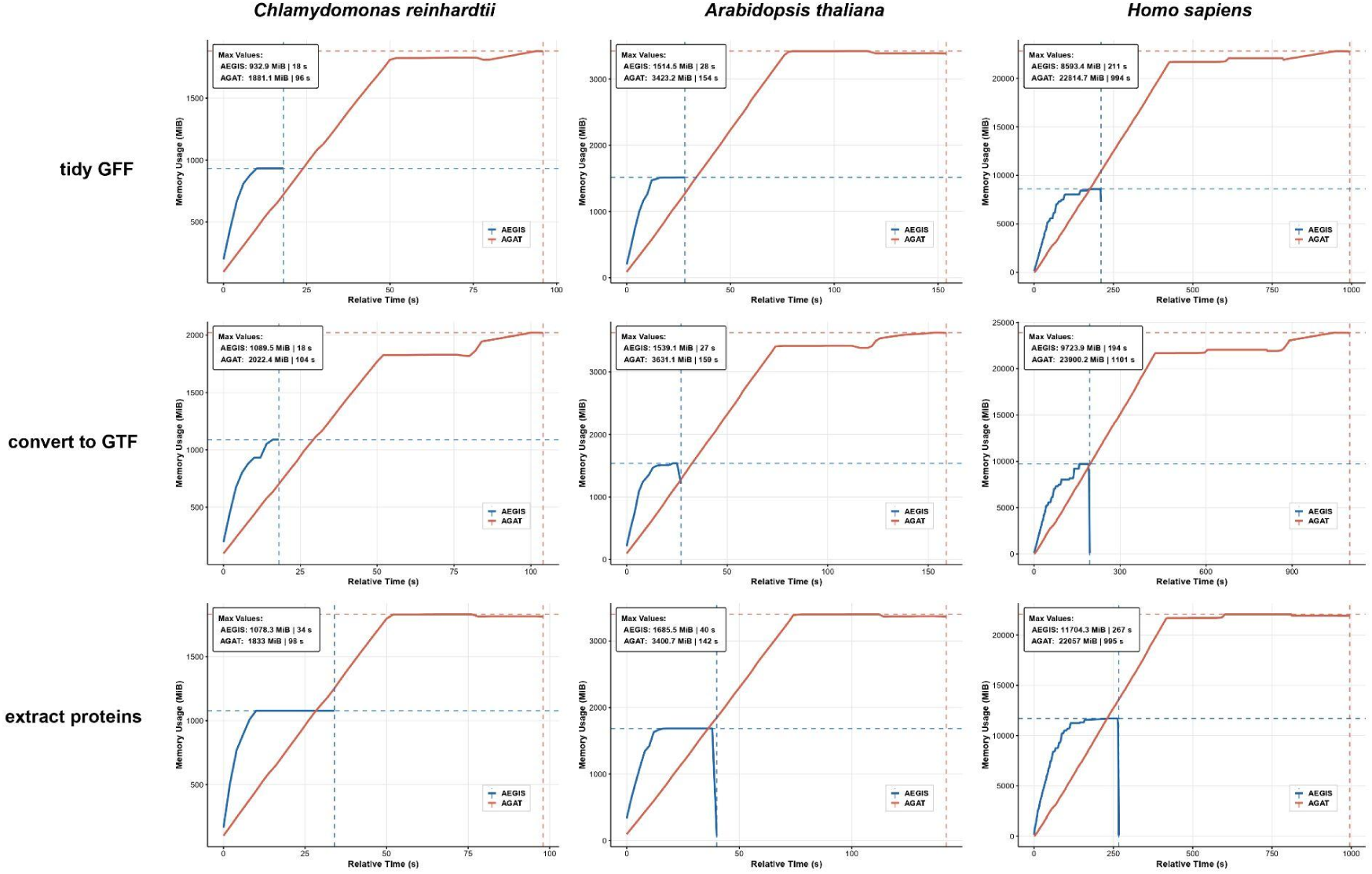
Computational performance benchmark of AEGIS and AGAT. Memory usage (MB) and relative execution time (s) were compared across three representative genomes: *Chlamydomonas reinhardtii* (small), *Arabidopsis thaliana* (medium), and *Homo sapiens* (large). Performance was evaluated for three core tasks: (top) GFF tidying (AEGIS ***tidy*** vs. AGAT agat_convert_sp_gxf2gxf.pl), (middle) GTF conversion (AEGIS ***reformat*** vs. AGAT agat_convert_sp_gff2gtf.pl), and (bottom) protein extraction (AEGIS ***extract*** vs. AGAT agat_sp_extract_sequences.pl).

